# Context-Dependent PSIICOS: A Novel Framework for Functional Connectivity Estimation Accounting for Task-Related Power Leakage

**DOI:** 10.1101/2024.12.30.630771

**Authors:** Daria Kleeva, Alexei Ossadtchi

## Abstract

Functional connectivity (FC) analysis using non-invasive neuroimaging methods, such as MEG and EEG, is often confounded by artifacts from spatial leakage and task-related power modulations. To address these limitations, we present Context-Dependent PSIICOS (CD-PSIICOS), a novel framework that improves the estimation of FC by incorporating task-specific cortical power distributions into the projection operator applied to the vectorized sensor-space cross-spectrum. Unlike the original PSIICOS (Phase Shift Invariant Imaging of Coherent Sources) approach, designed to suppress spatial leakage from all the sources, CD-PSIICOS dynamically adjusts the projection based on the active source distribution, enabling more accurate suppression of spatial leakage while preserving true zero-phase interactions. We validated CD-PSIICOS using realistic simulations and a multi-subject MEG dataset. The results demonstrate that CD-PSIICOS outperforms the original PSIICOS in suppressing artifacts at the lower projection ranks, maintaining robust detection of functional networks across theta and gamma frequency bands. By requiring lower projection ranks for optimal performance, CD-PSIICOS facilitates the reconstruction of physiologically relevant networks with improved sensitivity and stability.

## 1. Introduction

Existing evidence strongly indicates that healthy brain functioning and its dynamical properties rely on connectivity between cell assemblies at various resolution levels, which reflects communication between different brain areas (Greicius, 2008). The task of mapping the human connectome remains one of the most significant challenges in neuroimaging (Bassett and Bullmore, 2009), limiting our understanding of the fundamental principles underlying brain function.

Noninvasive time-resolved neuroimaging techniques such as EEG and MEG, combined with source localization methods, offer the potential to examine large-scale functional connectivity (FC) in terms of space, time, frequency, and phase-lag (Ossadtchi et al., 2018). These unique benefits arise from measuring the brain’s electrical activity with sensor arrays covering the entire head.

Traditional approaches to functional connectivity in MEG and EEG have largely relied on linear measures sensitive to frequency-specific phase and amplitude relationships across brain regions (Marshall et al., 2002). They operationalize the communication through coherence hypothesis Fries (2015); Varela et al. (2001). These methods assume narrowband oscillatory coupling and are effective in detecting synchronous rhythms, but they have important limitations — particularly in the context of aperiodic or nonlinear neural interactions. Recent studies have emphasized the need to move beyond the traditional frameworks. For instance, measures based on temporal irreversibility — such as lagged amplitude envelope correlations — may outperform classical functional connectivity metrics in distinguishing cognitive states (Tewarie et al., 2023), and state-space models have been proposed to disentangle periodic and aperiodic signal components in a principled, data-driven manner (Matsuda and Komaki, 2017) that can inspire real-time phase-tracking frameworks robust to 1/f background activity (Fedosov et al., 2024) or, conversely, characterize coupling in the absence of sustained rhythms. Furthermore, alternative classes of connectivity measures, such as transfer entropy (Wibral et al., 2014), have been proposed to capture directed and nonlinear interactions that are not accessible to coherence-based approaches. While our current method focuses on linear, phase-based coupling in the frequency domain, we acknowledge these broader developments and view them as complementary to the framework presented here.

To effectively quantify frequency-specific phase coupling, it is essential to employ such measures as cross-spectrum, coherence and their derivatives such as PLV, PLI and others (Bastos and Schoffelen, 2016; Nolte et al., 2004, 2008; Stam et al., 2007), which facilitate the detection of consistent phase relationships over time. Based on the “communication through coherence” principle (Fries, 2015) tracking the dynamics of phase synchrony between the activity of neuronal assemblies allows for exploring the dynamic processes of inter-regional communications implemented by the brain.

However, such FC measures computed from the non-invasively measured electromagnetic signals are prone to Type I and Type II errors. Firstly, the artifacts of volume conduction, or spatial leakage (SL) in MEG, often result in spurious connections between brain regions that do not reflect true neuronal interactions. This problem was addressed with various methods. The most potent approach that gave rise to a family of methods for tracking phase synchrony is based on exploring the imaginary part of the sensor-space coherence (Nolte et al., 2004). The imaginary part of coherence is devoid of the volume conduction contributions due to the instantaneous nature of the latter (Hamalainen et al., 1993) which furnishes certain control over the false positives in the non-invasive FC studies. However, the methods relying on the imaginary part of coherence struggle to detect the networks whose nodes exhibit zero or close to zero mutual phase-lags Ossadtchi et al. (2018) and are therefore prone to generating false negatives. Ironically, such instantaneous coupling scenarios are ubiquitous which has been shown both theoretically Chawla et al. (2001); Rajagovindan and Ding (2008); Fischer et al. (2006); Vicente et al. (2008); Esfahani and Valizadeh (2014); Gollo et al. (2014) and empirically (Gray et al., 1989; Frien et al., 1994; Roelfsema et al., 1997; Rodriguez et al., 1999; Fell et al., 2001; Havenith et al., 2009; König et al., 1995) in numerous studies. In particular, the evidence for genuine zero-phase interactions comes from spike-train recordings, which revealed that most functional connections between homotopic interhemispheric regions operate with zero-phase delay (Engel et al., 1991). Similarly, invasive recordings in humans have shown near-zero phase relationships, with delays significantly shorter than those predicted by axonal conduction alone (Uhlhaas et al., 2009). Several plausible mechanisms may underlie these observations (Rajagovindan and Ding, 2008; Gollo et al., 2014; Fischer et al., 2006). One possibility is bidirectional communication, where mutual coupling between regions leads to stable phase alignment. Alternatively, common input from a third structure can synchronize the dynamics of two otherwise unconnected areas into a near-zero-lag regime. A more complex explanation is dynamical relaying, wherein the activity of two distinct neuronal groups is relayed via the third neuronal population, causing the dynamics to be redistributed and self-organized zero-phase coupling to emerge. These findings suggest that instantaneous or near-instantaneous synchronization is not only theoretically tractable but also physiologically grounded.

In (Ossadtchi et al., 2018; Altukhov et al., 2023) we introduced PSIICOS (Phase shift invariant imaging of coherent sources) approach, which, by operating on the vectorized M *×* M sensor-space cross-spectrum, effectively suppresses the spatial leakage and without completely removing the genuine true zero-phase coupling.

PSIICOS uses a specifically designed projection operation that yields significant suppression of the volume conduction effect in the real part of the cross-spectrum while retaining the contributions modulated by the off-diagonal elements of the source-space cross-spectral matrix. However, due to the fundamental reasons, it is impossible to completely zero the spatial leakage contributions and overly powerful task-related induced activity generated by the genuinely uncoupled cortical sources may still lead to the false positives (Schoffelen and Gross, 2009; Bastos and Schoffelen, 2016; Kleeva and Ossadtchi, 2021). In these cases the power and selectivity of PSIICOS projection may appear insufficient due to its universality as it is designed to suppress the field spread contributions and the associated mutual spatial leakage from *all* cortical sources (Altukhov et al., 2023).

In this paper we extend the original PSIICOS technique by constructing the projection operator that takes into account the distribution of active sources pertinent to the analyzed data. Instead of building the projection operator using the *M* ^2^ *×* 1 auto 2-topographies of *all* sources present in the forward model matrix here we propose to take into account the cortical power distribution context.

After describing the methodological aspects of the context-dependent PSIICOS (CD-PSIICOS) technique we first validate this framework through testing using the realistically simulated data. We show its superior performance as compared to the generic PSIICOS and explore the extent to which the CD-PSIICOS depends on the particular inverse operator used to obtain the distribution of the induced cortical activity power. We also show that CD-PSIICOS requires a lower optimal projection rank as compared to the original PSIICOS which means that potentially more of the genuine close-to-zero-phase networks will be retained after the projection.

Then, we demonstrate the application of CD-PSIICOS to real data using a multi-subject 275-channel CTF MEG dataset recorded from nine volunteers during an audio-motor integration task.

We explore the stability of the observed networks and the extent to which they are affected by the variation of the PSIICOS projection rank. We report the spatial and the temporal profiles of the discovered networks in the alpha, beta, theta and gamma bands and assess their physiological validity given the behavioral paradigm used to collect the data.

We also verify that the discovered networks indeed correspond to the induced activity with random absolute phase and the persistent phase difference that appeared to be close to zero for the networks derived from the CD-PSIICOS de-biased real part of the cross-spectrum.

## 2. Methods

### 2.1 Generative model

The EEG/MEG setup captures electrophysiological data through an M-channel device that senses the activity of neuronal sources distributed over the cortical surface. This surface is digitized into a mesh with *N* vertices to comprise the source space according to the brain’s anatomical features. Each vertex in the mesh is linked to a neuronal source that is approximated by an equivalent current dipole (ECD). The data from K trials is produced by mapping ECD activity to sensor measurements, which mixes the activity from different sources on each sensor and contaminates the mixture with the additive measurement noise. Taking into consideration only induced activity, typically analyzed in the time-frequency domain, we can describe the observed data at a given time *t*, frequency *f* within a trial *k* using the following generative equation:

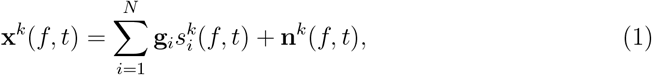

where **g**_*i*_, the *i*-th column of forward operator **G**, represents the topography of the *i*-th source, 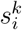 represents the *i*-th source activity at trial *k* and **n**^*k*^(*f, t*) models zero mean spatially uncorrelated additive sensor noise.

Based on the theory introduced in (Ossadtchi et al., 2018; Altukhov et al., 2023), the task of FC estimation can be performed in one step without separate reconstruction of source activations 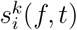.

To show this, we first substitute (1) in the definition of sensor-space cross-spectrum **C**^*xx*^(*f, t*) = *E*{**x**(*f, t*)**x**^*H*^ (*f, t*)} and obtain its vectorized form vec(**C**^*xx*^(*f, t*)) as

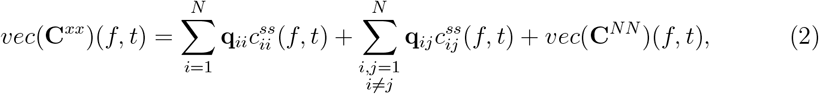

where 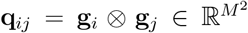 are the *M* ^2^ *×* 1 2-topography vectors computed as the vectorized outer products of the *i*-th and the *j*-th columns **g**_*i*_ and **g**_*j*_ of the traditional forward model matrix **G**. This representation allows us to express the sensor-space cross-spectrum as a linear combination of such 2-topography vectors. For (*i ≠ j*) we will refer to **q**_*ij*_ = **g**_*i*_ ⊗ **g**_*j*_ as the interaction topography of the (*i, j*)-th pair and when *i* = *j* we will call this the auto 2-topography of the *i*-th source. As detailed in (Ossadtchi et al., 2018) these auto 2-topographies span the spatial leakage (SL) subspace. It is possible to suppress the spatial leakage using a projection operator and then focus on estimating 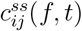 by formulating this task as a regression problem in the *M* ^2^-dimensional space.

### 2.2 Spatial leakage suppression and source space cross-spectrum estimation

As demonstrated in (Altukhov et al., 2023) given a set of symmetric auto 2-topographies

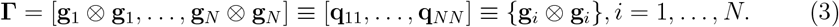

and the orthogonal projector **P** = **I** − **ΓΓ**^*†*^, where *†* denotes the pseudo-inverse, vector

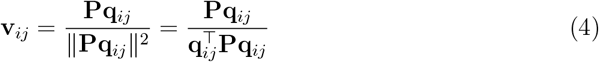

is the optimal filter that minimizes the cross-talk originating from the mutual spatial leakage when applied to the vectorized cross-spectral matrix (2) to estimate the cross-spectral coefficient 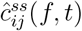 of the (*i, j*)-th source pair as

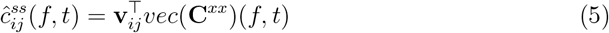

This expression forms the basis for the earlier proposed PSIICOS technique (Ossadtchi et al., 2018) that constructs a reduced rank PSIICOS projector matrix **P** using the first *R* left singular vectors of matrix **Γ**. These *R* left singular vectors are arranged as columns of matrix **U**_*R*_ and the rank-reduced projection operator is computed as 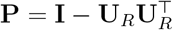 and is then applied to *vec*(**C**^*xx*^)(*f, t*) in order to suppress the undesired SL component, see (2).

Within the CD-PSIICOS approach introduced in this paper, we propose modifying the filtering process and to account for the distribution of source power in the particular data, namely the component 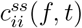 from equation (2). This can be achieved by weighting each column *i* of **Γ** by the estimated power of the *i*-th source prior to computing the SVD. This approach is detailed next.

### 2.3 CD-PSIICOS pipeline

The standard processing procedure involves bandpass filtering the epoched data within the frequency band of interest. After filtering, we compute the outer products of the Hilbert transforms of this data for each epoch. By averaging these outer products across epochs, we obtain the time-varying sensor-space cross-spectral coefficients, denoted as **C**^*xx*^(*f, t*). Finally, we estimate the averaged band-specific power distribution across the cortex using one of the standard linear inverse operators, **W**:

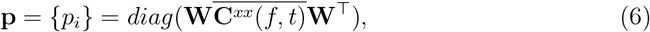

where the bar over the sensor-space cross-spectral matrix **C**^*xx*^(*f, t*) denotes its mean value computed over a specific frequency and time range of interest.

The main idea of CD-PSIICOS is to use the estimated source power to scale the contributions to the SL subspace matrix **Γ**. First, for simplicity and consistency with (3) we illustrate this for the 1-D case assuming fixed source orientations. Within the CD-PSIICOS we then construct the modified SL matrix **Γ**^*CD*^ using auto 2-topographies scaled with the corresponding power estimate as its columns:

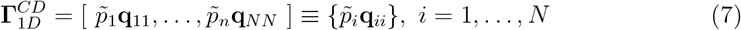

Here 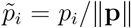 is the normalized frequency band and time interval specific power of the *i*-th source, *i* = 1, … , *N* .

While the assumption regarding the orientation of source dipoles orthogonally to the cortical surface is plausible (Okada, 1983), accurately applying this orientation constraint requires extremely dense cortical meshes. Given the combinatorial nature of the pairwise interaction modeling, we instead use a moderately sparse source space with free orientations within the 2-D planes defined by the two dominant singular vectors of the gain matrices at each cortical location. Therefore, in our actual implementation instead of this 1D SL matrix 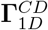 we use the matrix **Γ**^*CD*^ accounting for spatial leakage emanating from sources with arbitrary orientations in the locally tangential planes. To form this matrix we use the observation described in the original PSIICOS paper (Ossadtchi et al., 2018) that the SL subspace of a source at the *i*-th location is spanned by the three 2-topography vectors 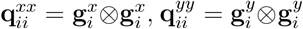 and 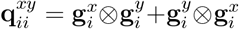 where 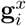 and 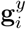 are the topographies of dipoles oriented along the *x* and *y* directions in the locally tangential plane at the *i*-th cortical location. Therefore, we form **Γ**^*CD*^ by stacking these triplets of column vectors weighed by the corresponding normalized source power at each *i*-th location as:

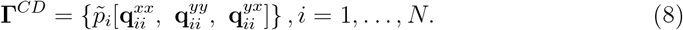

The associated PSIICOS projection matrix **P**^*CD*^ is then computed as

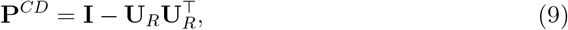

where **U**_**R**_ is the matrix of the first *R* left singular vectors of the power distribution context dependent **Γ**^*CD*^ defined above. Projection rank *R* corresponds to the number of the left singular vectors used to construct the projection operator.

The rest of the pipeline remains the same as in the original PSIICOS. To estimate the source space cross-spectral coefficient 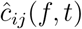 we simply regress the vectorized cross-spectrum to the projected and appropriately normalized 2-topography vector 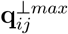 corresponding to the interacting pair (*i, j*) oriented to maximize 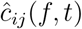:

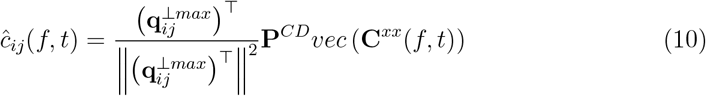

To implement this, we scan the unvectorized projected cross-spectrum for the interaction between the *i*-th and the *j*-th sources using the pair of topographies of two orthogonal locally tangential dipoles in each of the two source locations. This results in a 2×2 cross-spectral matrix per source pair. Then, similarly to the DICS technique (Groß et al., 2001) we use singular value decomposition (SVD) of this matrix to extract the pair of orientation vectors that maximizes the coupling strength. This procedure ensures that the interaction estimates 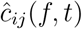 are not limited to fixed source orientations and remain sensitive to the dominant coupling directions in the source space.

Note that 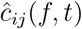 is unnormalized, meaning we do not divide this cross-spectral coefficient by the product of spectral power values at each node. We avoid this step as our earlier experiments showed that such a normalization adversely affects the sensitivity to source pairs coupled at zero- or close-to-zero phase lags.

In the original PSIICOS the only adjustable parameter of the projection was its rank *R*, which is selected based on the optimal trade-off between the suppression of the SL subspace and the preservation of the real part of the cross-spectrum subspaces. In this study, we specifically examined how the CD-PSIICOS projection behaves at different ranks in comparison to the original projection. Furthermore, since the CD-PSIICOS projector is influenced by the choice of the inverse operator, its regularization parameters, and weighting techniques, we also investigated the robustness of the observed solutions with respect to these parameters.

### 2.4 Realistic simulations

To evaluate the proposed approach, we conducted a series of realistic Monte Carlo (MC) simulations inspired by those used in the original paper (Ossadtchi et al., 2018). We utilized a cortical surface model containing 15 000 vertices and the corresponding high-resolution forward-model matrix. For each cortical location the forward matrix had 2 topographies representing the sources in the locally tangential plane. All simulations were performed using the 204 planar gradiometers of the Elekta Neuromag system to match the sensor configuration used in the previous PSIICOS studies (Ossadtchi et al., 2018; Altukhov et al., 2023).

Each dataset from MC simulations included 100 epochs. To model the induced activity of coupled sources, we used two 10 Hz harmonics with random phases at the start of each epoch. Phase jitter was drawn from a random distribution within the range of [−*π/*20, *π/*20], with a mean phase lag of *ϕ* = *π/*20. In our study, we simulated two truly coupled networks and one network characterized by a lack of coupling (phase jitter assigned as 2*π*) and excess power (source activation for the network scaled by a factor of 3). This excess power was expected to result in high cross-spectrum values and overestimation of the FC. For each simulation run, the locations of the true coupled and uncoupled networks were randomly varied. We considered two SNR conditions: 0.5 and 1.0.

Brain noise was modeled as the sum of narrow-band signals drawn from a Gaussian distribution. These signals were filtered using a 5th-order band-pass IIR filter, corresponding to the theta (4-7 Hz), alpha (8-12 Hz), beta (15-30 Hz), and gamma (30-50 Hz, 50-70 Hz) frequency ranges. The signal-to-noise ratio was defined as the ratio of the Frobenius norms of the induced and brain noise time series filtered in the band of interest (8-12 Hz). The power spectral density matched the spectral characteristics of EEG or MEG data. At each trial these time series we assigned to a thousand of randomly picked source locations that were reselected for each trial. To impose the spatial correlation structure and to obtain the brain-noise data in the sensor space we projected the sources time series into the sensor space through a linear combination of the forward matrix columns corresponding to the locations of the randomly picked sources.

Prior to estimating the sensor-space cross-spectrum and constructing the PSIICOS projector, we used a 10-fold sparser cortical model with 1503 nodes downsampled from the original one. This reduction also serves to mitigate the ”inverse crime” in simulation studies — that is, to prevent artificially optimistic results due to the reuse of the same forward model for both signal generation and reconstruction. By intentionally introducing a mismatch between the dense source model used for simulation and the coarser model used for analysis, we mimic the real-world condition where the true biophysical model of the head is never precisely known. This strategy introduces a more realistic level of modeling uncertainty and better reflects the challenges faced in real-data applications.

In all simulations, we used the sensor-weighted overlapping-sphere (OS) head model (Huang et al., 1999) for forward modeling. This approach fits a sphere to the local head geometry on a per-sensor basis, allowing a rapid yet accurate approximation of field propagation. Our use of the OS model allows for extensive exploration of source interaction scenarios without incurring the computational burden of the fully-fledged BEM simulations. In contrast, for further analysis of the real MEG data we used a traditional BEM model aiming to represent the biophysics of the head as a volume conductor.

To evaluate the performance of CD-PSIICOS under different inverse modeling assumptions, the power of the sources incorporated into the CD-PSIICOS projector, see equation (8), was estimated using the standard inverse operators (MNE (Hämaläinen and Ilmoniemi, 1994) and sLORETA (Pascual-Marqui et al., 2002)) and the DICS frequency domain beamforming approach (Groß et al., 2001). The inverse operator **W** was applied to the original (non-projected with PSIICOS) sensor-space cross-spectrum averaged within the time-frequency patch as 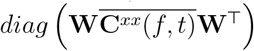 . Additionally, as the ultimately best scalar (albeit unavailable in practice) we performed simulation experiments with auto-topographies weighted with respect to the ground-truth active sources. To this end, the auto-topographies corresponding to the ground-truth active sources were multiplied by the weighting coefficient, while all other topographies remained unchanged. For the majority of the reported results the weighting coefficient for the ground-truth solution was set to 10. We examined the influence of the weighting coefficient parameter in a set of separate experiments reported below.

Some properties of the solutions were additionally analyzed in a setting illustrated in Figure 1. Here we are presenting the spatial (A) and temporal (B) profiles of the modeled networks we used as the ground truth in one of our experiments. Here we simulated 2 bilateral coupled and one uncoupled pair of sources. The temporal profiles of the coupling strength shown in Figure 1.B illustrate the partially overlapping activity of the two true networks. The synchrony profile of the third pair of bilateral nodes is a flat line indicating no true coupling present over the considered time interval.

**Figure 1:**
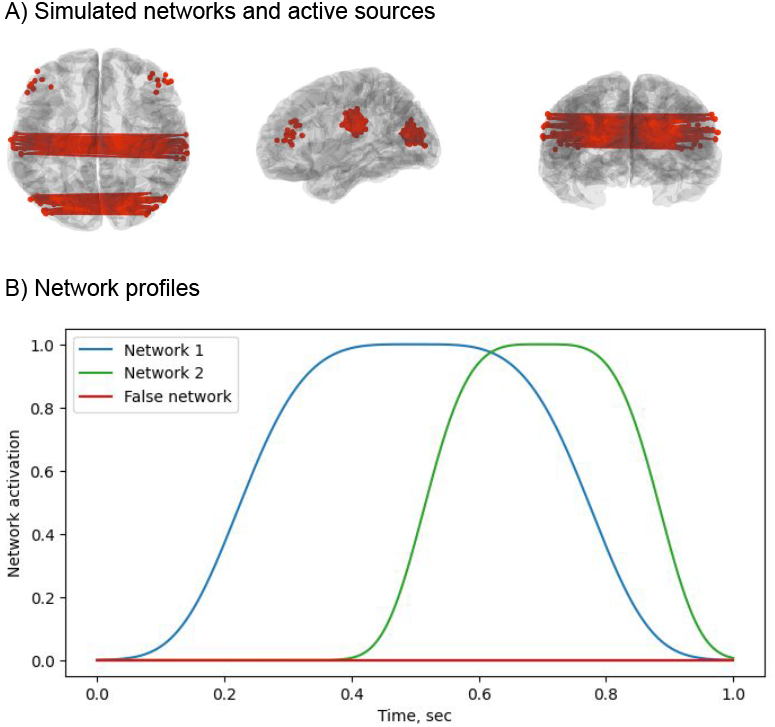
Modeled networks. The spatial (A) and the temporal (B) profiles of the simulated networks, a representative example.

To evaluate the networks detection accuracy in our experiments we use Receiver Operating Characteristics (ROC) illustrating the achievable trade-off between the sensitivity and specificity characteristics of the detector. To this end we estimate the source-space cross-spectral coefficients 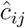 for all source pairs and apply a sweeping threshold over the range of the obtained values. Starting from min(*c*_*ij*_) and increasing to max(*c*_*ij*_), we compute true positives (TP), false positives (FP), true negatives (TN), and false negatives (FN) at each threshold. A source pair is considered a detected interaction if its score exceeds the threshold. The resulting ROC curve plots the true positive rate against the false positive rate across all thresholds, providing a threshold-independent measure of discriminative performance. As a scalar summarizing the attainable discriminability we used the area under the ROC curve metric, ROC AUC.

We used the standard definitions of the sensitivity and specificity metrics:

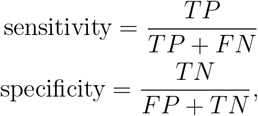

where *TP* , *FN* , *FP* and *TN* are the numbers of true positives, false negatives, false positives and true negatives correspondingly.

In our simulations, each truly coupled network corresponded to a pair of nodes in the high-resolution cortical mesh. For estimating source-space cross-spectral coefficients, we used a 10 times sparser cortical mesh to avoid the ”inverse crime”. A pair of sources was considered a true connection if their coordinates fell within *d* = 1.5 cm from the original coordinates of the coupled sources.

It is important to note that truly coupled sources constitute only a small fraction of all possible connections. Consequently, a combination of the high true negative rate and low false positive rate is expected from a useful method.

### 2.5 Real data

We confirmed the suggested method’s effectiveness by employing the real MEG data for validation. The participants were drawn from an open dataset of MEG recordings (Nugent et al., 2022). As stated in the original paper, eligible participants were adults aged 18 or older, in good health, fluent in English, and capable of providing informed consent. The electronic consent for online screening and the written consent for the subsequent recordings were obtained from each of the participants. Exclusion criteria included significant health conditions, substance use, suicidal behavior, abnormal clinical findings, or low educational attainment or cognitive ability. The dataset was publicly released as part of the NIH BRAIN Initiative. The data were collected under the NIMH Intramural Research Protocol titled “Recruitment and Characterization of Healthy Research Volunteers for NIMH Intramural Studies”, registered at ClinicalTrials.gov under ID NCT033046. The study procedures were approved by the Institutional Review Board of the National Institute of Mental Health.

A total of nine healthy adult individuals (average age: 29.7, 4 males, 5 females) volunteered to take part in the study. MEG data were collected using a 275-channel CTF MEG system (CTF MEG, Coquiltam BC, Canada). To correct for noise, third-order gradient balancing was employed. Data collection was conducted at the sampling rate of 1200 Hz, with a quarter-Nyquist filter set at 300 Hz.

During the experiment, the participants underwent a three-stimulus oddball task that involved presentation of the standard 1 kHz tone, the higher-pitched 1.5 kHz target tone and the white noise. Participants were asked to respond to the target tone by pressing a button.

The bandpass-filtered data underwent epoching within a window of -0.2 to 0.8 seconds relative to the stimulus onset, and the baseline correction was applied to the segment starting 200 ms before the stimulus onset. Subsequently, sensor-space cross-spectral time series were computed using the averaged outer products of Hilbert-transformed epochs. The PSIICOS projection variants were applied to this cross-spectral timeseries matrix. Inverse modeling was performed using the time-averaged cross-spectral data. The participant’s anatomical MRI was used with FreeSurfer software to reconstruct the cortical surface model. The number of vertices in the individual cortical meshes slightly varied but was kept around *N* = 1500. The boundary element model (BEM) model was employed for forward computation using MNE Python (Gramfort et al., 2013).

To obtain the group-level results we used the ’fsaverage’ cortical source space with 1284 sources to which we morphed individual source spaces. The network detection results of each individual subject were morphed to the ‘fsaverage’ space for the group-level averaging.

## 3. Results

### 3.1 Realistic simulations

The results of the Monte Carlo simulations were consistent across both SNR levels (0.5 and 1), indicating the robustness of the CD-PSIICOS framework to moderate changes in the signal strength. The results indicate that CD-PSIICOS approach outperforms the standard one in terms of ROC AUCs, particularly at lower projection ranks (see Fig. 2, 1.A-2.A). Specifically, for SNR=0.5, at a fixed projection rank of 40 (Fig. 2, 1.B), the CD-PSIICOS with MNE-derived weighting demonstrates superior performance compared to both solutions based on sLORETA and DICS, and shows slightly better performance even compared to the ground-truth-weighted solution. The gap between CD-PSIICOS with MNE and the ground truth variant – along with the other CD-PSIICOS versions – becomes more pronounced at the projection rank 80. The standard PSIICOS solution lags significantly behind, showcasing reduced sensitivity and specificity at this lower rank. At the higher ranks, the overall performance of all CD-PSIICOS approaches converges, while standard PSIICOS shows significant improvement at this rank but still fails to match the accuracy and specificity achieved by the CD-PSIICOS solutions, particularly in the lower FPR range. Finally, at a high projection rank of 500 (Fig. 2, 1.B-2.B), the differences between the methods diminish further, with all variants of CD-PSIICOS and the standard PSIICOS performing comparably. For SNR = 1, all CD-PSIICOS variants performed similarly across projection ranks, demonstrating consistent accuracy and robustness. Only at the projection rank of 40 (Fig. 2, 2.B), and specifically in the low-FPR range, did the MNE-weighted version show a slight advantage over the others. In contrast, the standard PSIICOS method continued to underperform in this regime, exhibiting noticeably lower sensitivity and reduced specificity compared to all CD-PSIICOS solutions.

**Figure 2:**
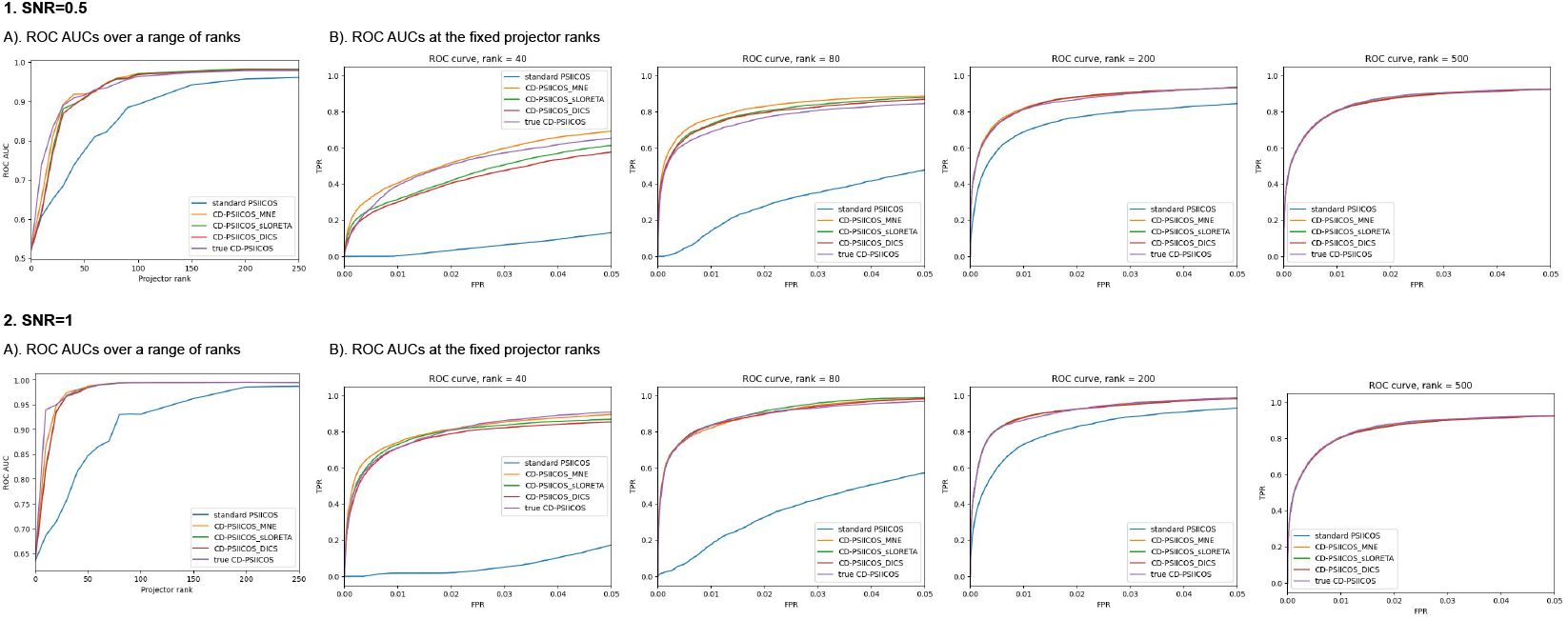
The impact of the standard PSIICOS approach and CD-PSIICOS variations on ROC AUCs across various projector ranks (A) and the detailed comparison of ROC curves at the fixed ranks (B)

To examine the properties of the obtained solutions using a representative example, we first obtained power profiles on the cortex using various inverse solutions. As evident from Fig. 3 (A-C), the activations of the sources align with the ground truth locations of the uncoupled, exceedingly active sources (Fig. 3 (D)). Notably, sLORETA and DICS solutions exhibit more diffuse patterns compared to the MNE.

**Figure 3:**
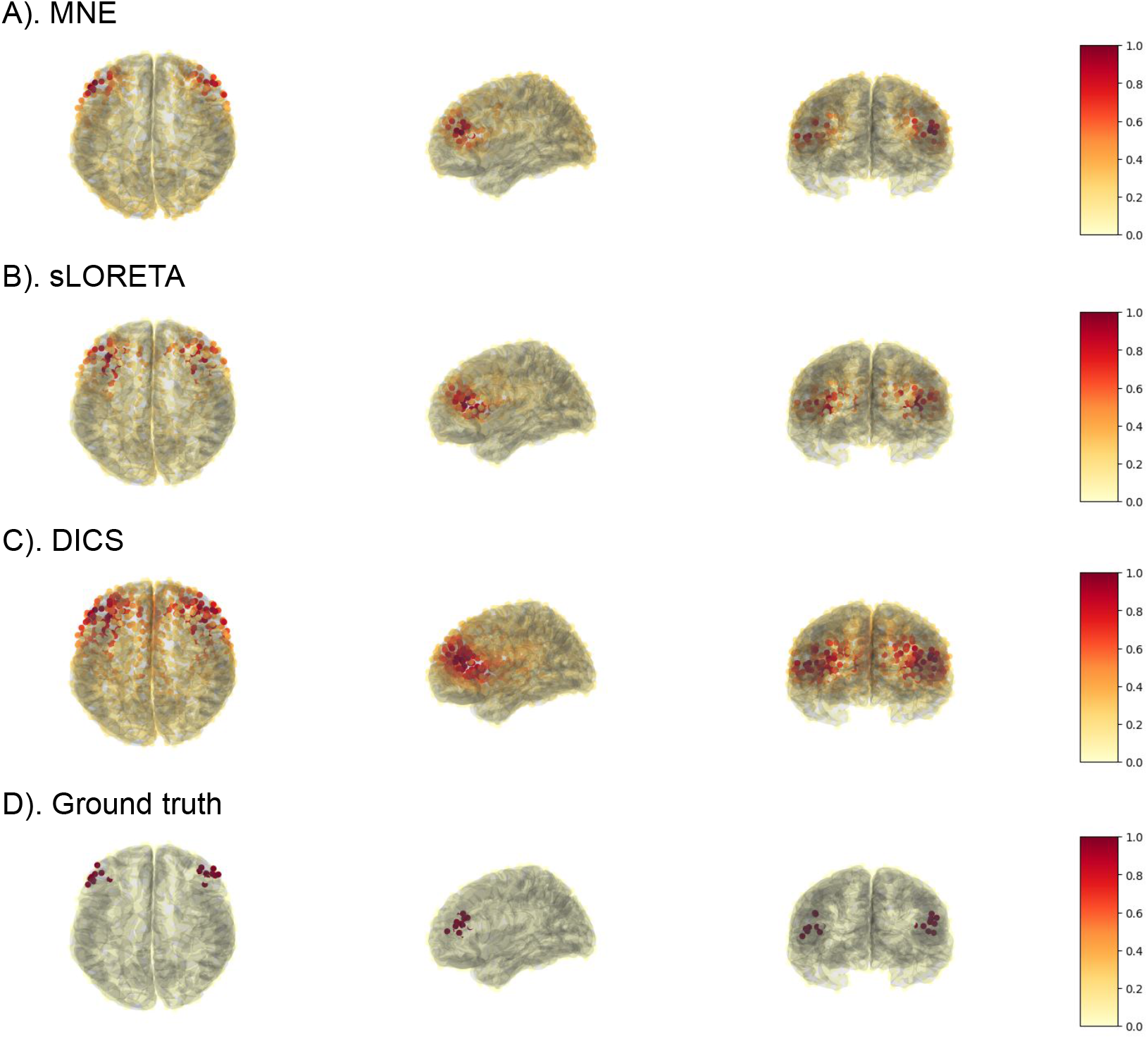
Power profiles obtained after applying inverse operators to the original sensor-space cross-spectrum: (A) MNE, (B) sLORETA, (C) DICS; and ground truth locations of active sources (D)

To evaluate the characteristics of the obtained PSIICOS and CD-PSIICOS projectors for the representative example, we present the singular value spectra showing the 100 largest singular values obtained for each set of 2-topographies (see Fig. 4). It is evident that the matrix of 2-topographies scaled by the corresponding source power values exhibits a much faster decay of singular values compared to the standard PSIICOS. This indicates that CD-PSIICOS can suppress the volume conduction contribution to the cross-spectrum more efficiently due to focusing on the active sources.

**Figure 4:**
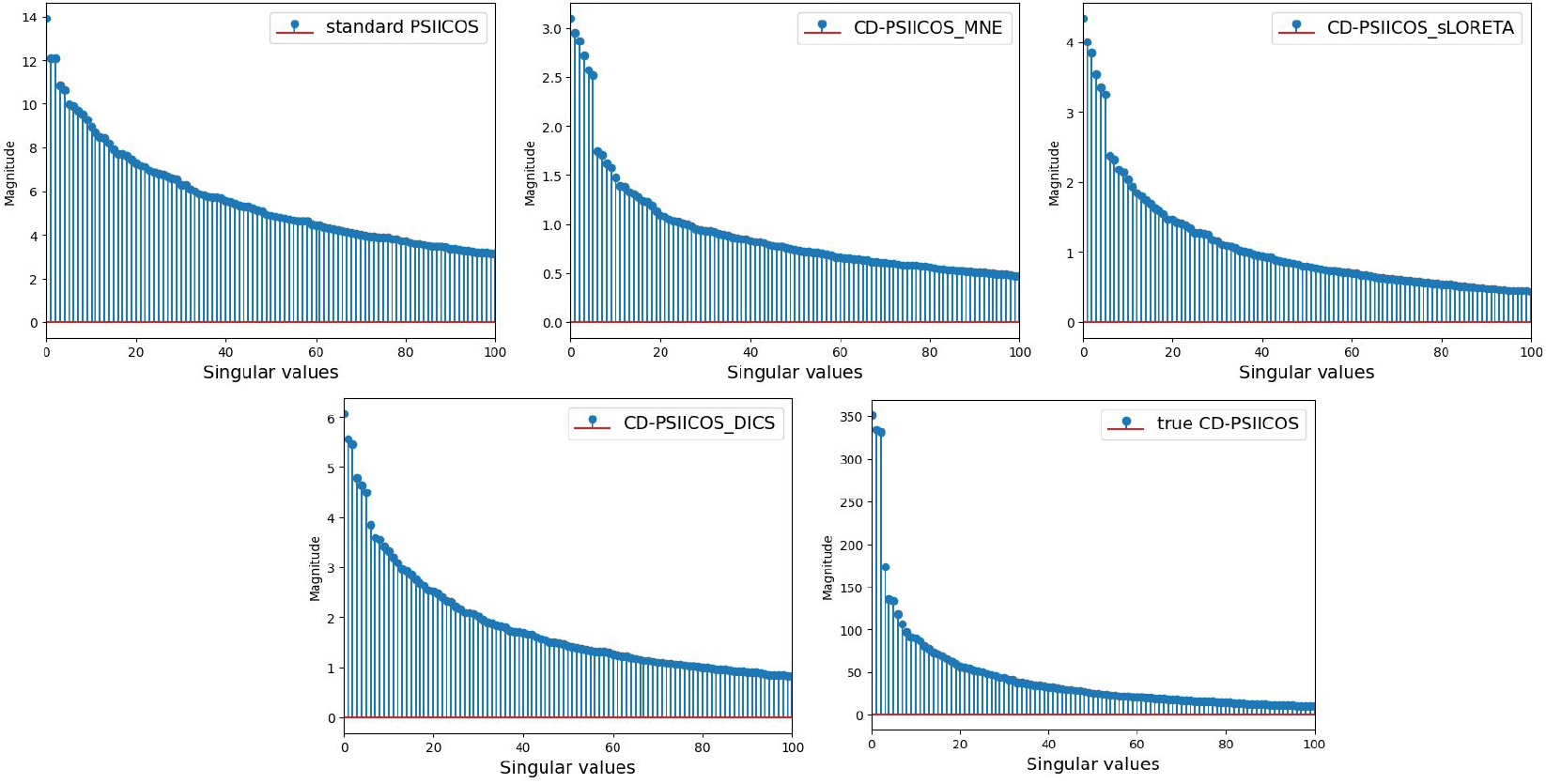
Singular values for each set of 2-topographies: (A) standard approach (A) CD-PSIICOS & MNE, (B) CD-PSIICOS & sLORETA, (C) CD-PSIICOS & DICS; CD-PSIICOS & and ground truth locations of active sources (D)

To enable consistent qualitative comparison between methods, we applied a fixed thresholding procedure across all conditions in the representative example: for each reconstructed network, the top *N* = 300 source pairs with the largest real-valued cross-spectral coefficients were selected and visualized. This approach ensures that the spatial patterns reflect the strongest estimated connections without introducing bias due to differing overall connection densities between methods.

The comparison of the standard PSIICOS approach and CD-PSIICOS variations (see Fig. 5) revealed that at low projector ranks (20, 40, 80), the standard PSIICOS was biased towards the clusters of active uncoupled frontal sources. Even at a higher projector rank of 200, while the true networks were reconstructed, residual false clusters from the active sources were still observed. On the other hand, the CD-PSIICOS approach based on the MNE, although showing false pairs due to remaining SL effects at rank=20, began to reconstruct the true networks from rank=40 onwards and was not biased towards the active uncoupled sources. CD-PSIICOS based on sLORETA, DICS, and the ground truth began to show consistent networks at the higher projector ranks.

**Figure 5:**
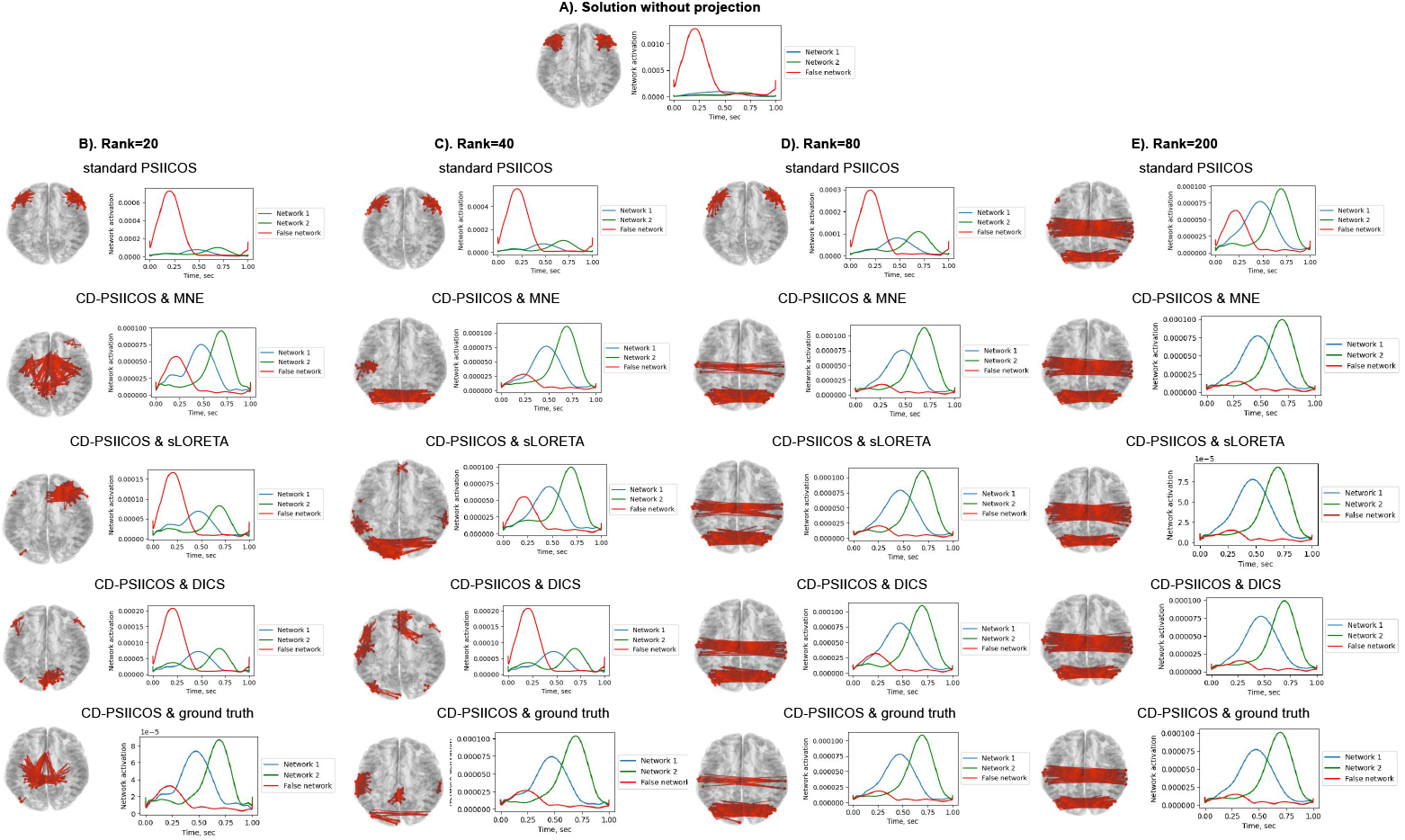
Spatial and temporal activation profiles of the networks reconstructed without PSIICOS approach (A) and with standard PSIICOS and CD-PSIICOS approaches at ranks equal to 20 (B), 40 (C), 80 (D) and 200 (E). For qualitative visualization, we applied a fixed-rank thresholding strategy: for each method, the top N=300 source pairs with the largest (real-valued) cross-spectral coefficients were selected and plotted as the estimated network. This uniform thresholding allows for side-by-side comparison of the spatial structure and leakage suppression capabilities of the methods under evaluation

To better understand the performance of CD-PSIICOS compared to the standard PSIICOS, we analyzed the Euclidean norms of the vectors comprising cross-spectral coefficients corresponding to the pairs of sources from the two groups: the truly coupled networks and the pairs from the regions with active but uncoupled sources (see Fig. 5). These norms represent the strength of connectivity estimated by the method, with higher values indicating stronger interactions. The idea is that a better method should yield a smaller norm of the cross-spectral coefficients for the falsely coupled pairs as compared to the truly coupled ones. For each group and each Monte Carlo trial, we compute a vector consisting of the real parts of the cross-spectral coefficients for all source pairs in that group. We then calculate the Euclidean norm of this vector to obtain a single scalar summarizing the aggregate interaction strength in the group. For the falsely coupled networks (solid lines), the CD-PSIICOS solutions show a much sharper reduction in the norm values compared to the standard PSIICOS solution (blue solid line), see Figure 6. This indicates that CD-PSIICOS is more effective at suppressing spurious connections caused by the spatial leakage, effectively improving the signal-to-noise ratio (SNR) for identifying genuine connectivity. The CD-PSIICOS variants showed a pronounced drop in the norm of the cross-spectral coefficients for uncoupled networks already at very low projection ranks — well below 50. The use of smaller projection ranks potentially leaves intact the contributions to the sensor-space cross-spectrum of the vast majority of the true zero-phase coupled pairs. Importantly, the norm of the cross-spectral coefficients vector for the falsely coupled networks in the CD-PSIICOS solutions fell below that of the truly coupled networks, further demonstrating the increased specificity of CD-PSIICOS. This explains the improved ROC performance observed for CD-PSIICOS, as the separation between true and false connectivity becomes more pronounced. By contrast, the standard PSIICOS approach achieves a comparable separation between true and false connections only at much higher projection ranks (starting at around 200). This illustrates the benefits of using the source power distribution context when building the CD-PSIICOS projector.

**Figure 6:**
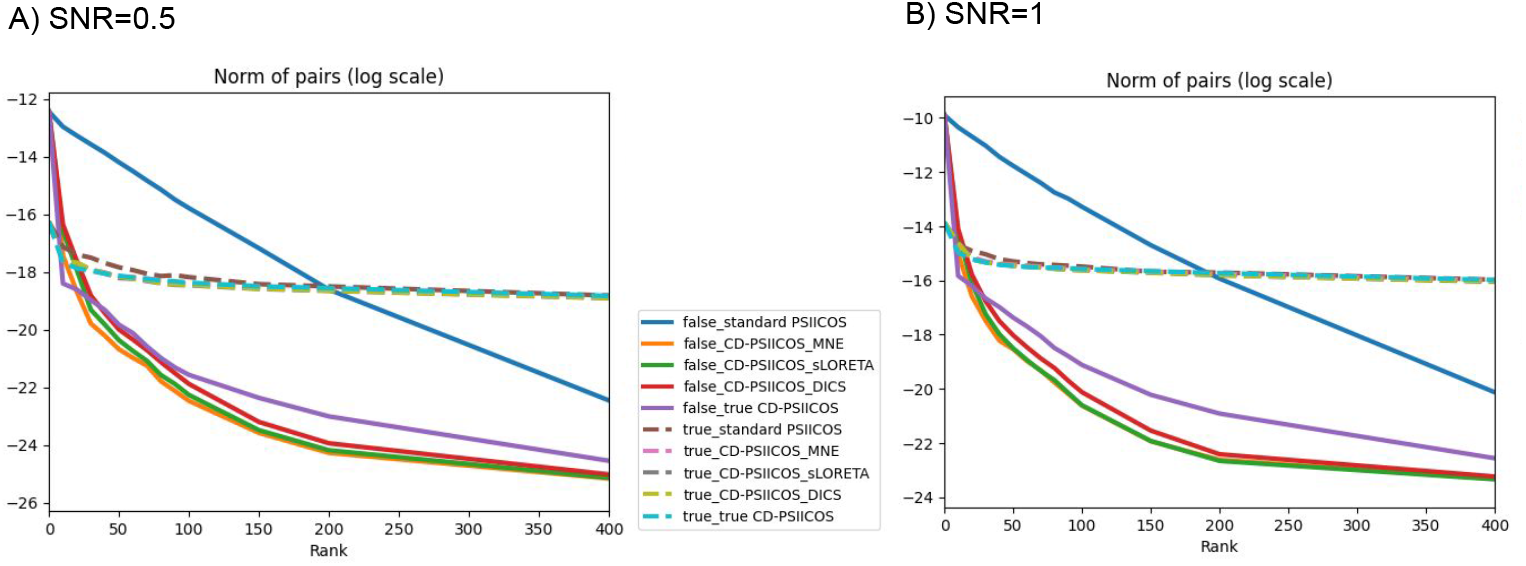
Norms of cross-spectral coefficients in the regions of truly coupled networks and uncoupled pairs depending on the type of solution and projection ranks for the simulations with SNR=0.5 (A) and SNR=1 (B)

To analyze the general characteristics of the proposed modifications, we computed attenuation curves similar to those in (Ossadtchi et al., 2018; Altukhov et al., 2023) that demonstrate the trade-off between suppressing spatial leakage and retaining the information about true interactions within source pairs. Originally, when the curves were computed exclusively using the model, we defined the spatial-leakage subspace and the subspace of the real part of the vectorized cross-spectrum using 2-topographies obtained from the forward model.

For a set of vectorized topographies (either auto-topographies representing spatial leakage, or 2-topographies representing the interactions), arranged into a matrix **Q**, we compute the attenuation at projection rank *R*, see equation (9), as the ratio of the Frobenius norms of the original and projected matrix **Q**:

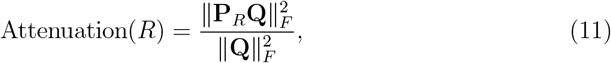

where **P**_*R*_ is the projection operator of rank *R* that removes the leading spatial leakage components. This value reflects the fraction of the original signal energy suppressed by the projector. To evaluate the trade-off between suppressing spatial leakage and preserving true interactions, we compute attenuation curves separately for auto-topographies and interaction topographies, and plot their difference:

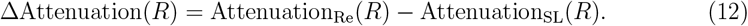

The peak of this differential curve highlights the projection rank where leakage is effectively removed while interaction-related patterns are preserved.

For the current comparison we computed the data-driven attenuation curves and accounted for a thresholded set of active sources with high power estimated through inverse modeling. We computed the norms of the auto-topographies (representing spatial leakage) and the 2-topographies (representing the interactions) for these sources and calculated the differences. The optimal rank is defined as the rank at which the attenuation curve reaches its maximum, representing the point where spatial leakage is sufficiently suppressed without compromising the interactions.

The obtained data-driven attenuation curves (see Fig. 7) reveal that CD-PSIICOS approaches achieve their optimal performance at significantly lower ranks compared to the standard PSIICOS. Specifically, for SNR=0.5 (see Fig. 7.A), the optimal rank is 500 for standard PSIICOS, 60 for CD-PSIICOS & MNE, 90 for CD-PSIICOS & DICS, and 100 for both CD-PSIICOS & sLORETA and CD-PSIICOS based on ground truth. For SNR=1 (see Fig. 7.B), the optimal rank is 500 for standard PSIICOS, 50 for CD-PSIICOS & MNE, 60 for both CD-PSIICOS & DICS and CD-PSIICOS & sLORETA , and 100 for CD-PSIICOS based on ground truth.

**Figure 7:**
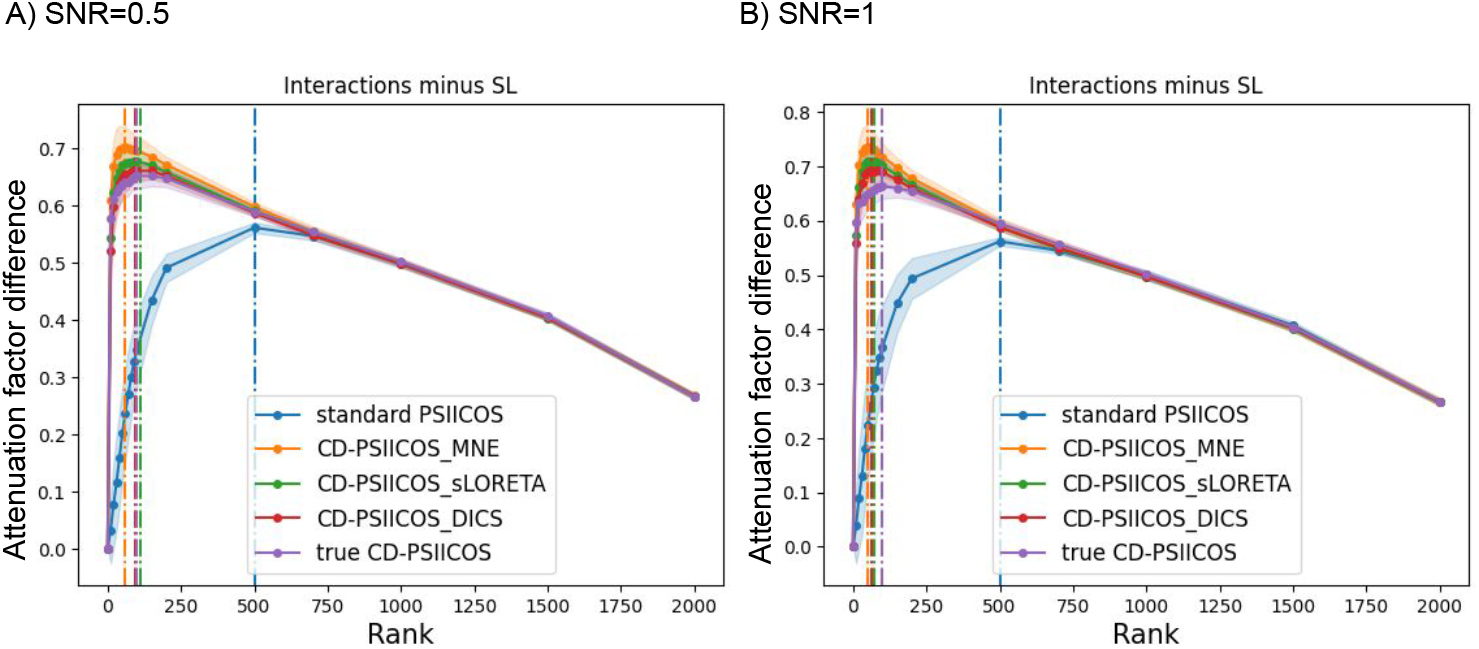
Attenuation difference curves (ΔAttenuation(*R*)) for the different ranks *R* of the PSIICOS and CD-PSIICOS projections applied to the simulated data with SNR=0.5 (A) and SNR=1 (B). The dashed lines represent the optimal projection ranks

Several variable parameters within the proposed approach may potentially affect the algorithm’s performance. These parameters include a) whether the thresholding is applied to inverse solutions coefficients 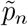 prior to weighting the corresponding auto-topographies in (8) or if the full inverse solution is utilized, b) the degree to which the sparse ground-truth-based coefficients are weighted, and c) the selection of the regularization parameter used in the inverse methods employed to estimate the source power distribution. Using CD-PSIICOS & MNE at rank=80, we demonstrated that the absence of thresholding resulted in higher ROC AUC compared to thresholding at other levels (see Fig. 8 , 1.A, 2.A, 3.A). For the CD-PSIICOS solution based on ground truth, we demonstrated that at a fixed rank of 80, the optimal coefficient is 10 in case when SNR=0.5 (see Fig. 8, 1.B). When SNR=1, the higher coefficients of 10 and 100 yield better performance (see Fig. 8, 2.B) Finally, the ROC curves based on the CD-PSIICOS solution with MNE demonstrated robustness across a wide range of regularization parameters. For SNR = 0.5 (see Fig. 8, 1.C), under-regularization (*λ* = 0 or 1*e*–3) led to a decline in ROC performance, but starting from *λ* = 1*e* − 1, the results stabilized. At SNR = 1 (see Fig. 8, 2.C), the differences across regularization values further diminished. In the representative example (see Fig. 8, 3.C), all tested regularization values resulted in comparable performance. The stability of the solution with respect to the regularization parameter is a critical aspect of the proposed approach, since regularization plays a pivotal role in inverse modeling, as it controls the trade-off between fitting the data and smoothing the solution to avoid overfitting to noise.

**Figure 8:**
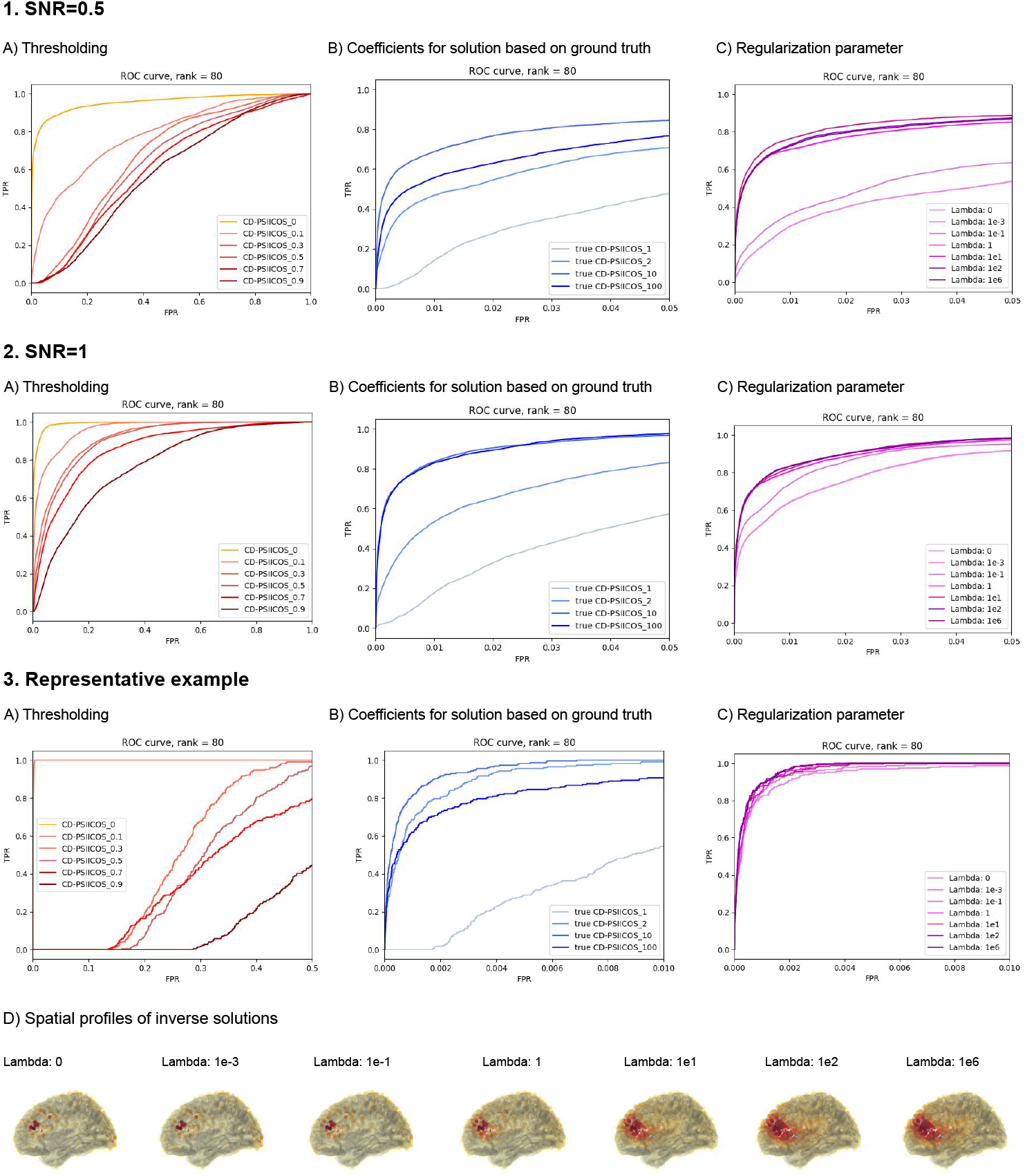
Dependence of performance on (A) thresholding, (B) degree of weighting and (C) regularization parameter with (D) spatial profiles of corresponding inverse solutions for the representative example. ”True CD-PSIICOS” refers to the variant where the projection operator was constructed using weighted auto-topographies based on the ground-truth knowledge of active (but uncoupled) sources. The condition “true CD-PSIICOS 1” corresponds to the standard PSIICOS solution, as no weightening is applied

The results from realistic simulations demonstrate that CD-PSIICOS significantly outperforms the standard PSIICOS approach, particularly at lower projection ranks, by effectively suppressing spatial leakage. Among the CD-PSIICOS variants, MNE-based weighting consistently showed superior performance, with optimal projection ranks significantly lower than those required by the standard approach. These findings confirm the robustness and efficiency of CD-PSIICOS in reconstructing functional networks, providing a solid foundation for its application to real MEG data.

While our simulations incorporate spectrally and spatially structured brain-like noise generated from large numbers of sources distributed over cortical surface, they still rely on Gaussian random processes and do not fully capture the non-Gaussian components present in real spontaneous electrophysiological activity. Incorporating more realistic noise models — including the non-Gaussian components remains an important direction for future work.

### 3.2 Real data

The grand average of the induced responses and the evoked activity revealed robust differences between the three experimental conditions (see Fig. 9 ). Specifically, in response to the target tones, we observed induced activity localized to frontal regions in the theta (4–7 Hz) and gamma (30–40 Hz) bands, peaking between 200 and 400 ms after the stimulus onset. Consequently, we focused our analysis on these frequency ranges within the 0–0.5-second time window post-stimulus.

**Figure 9:**
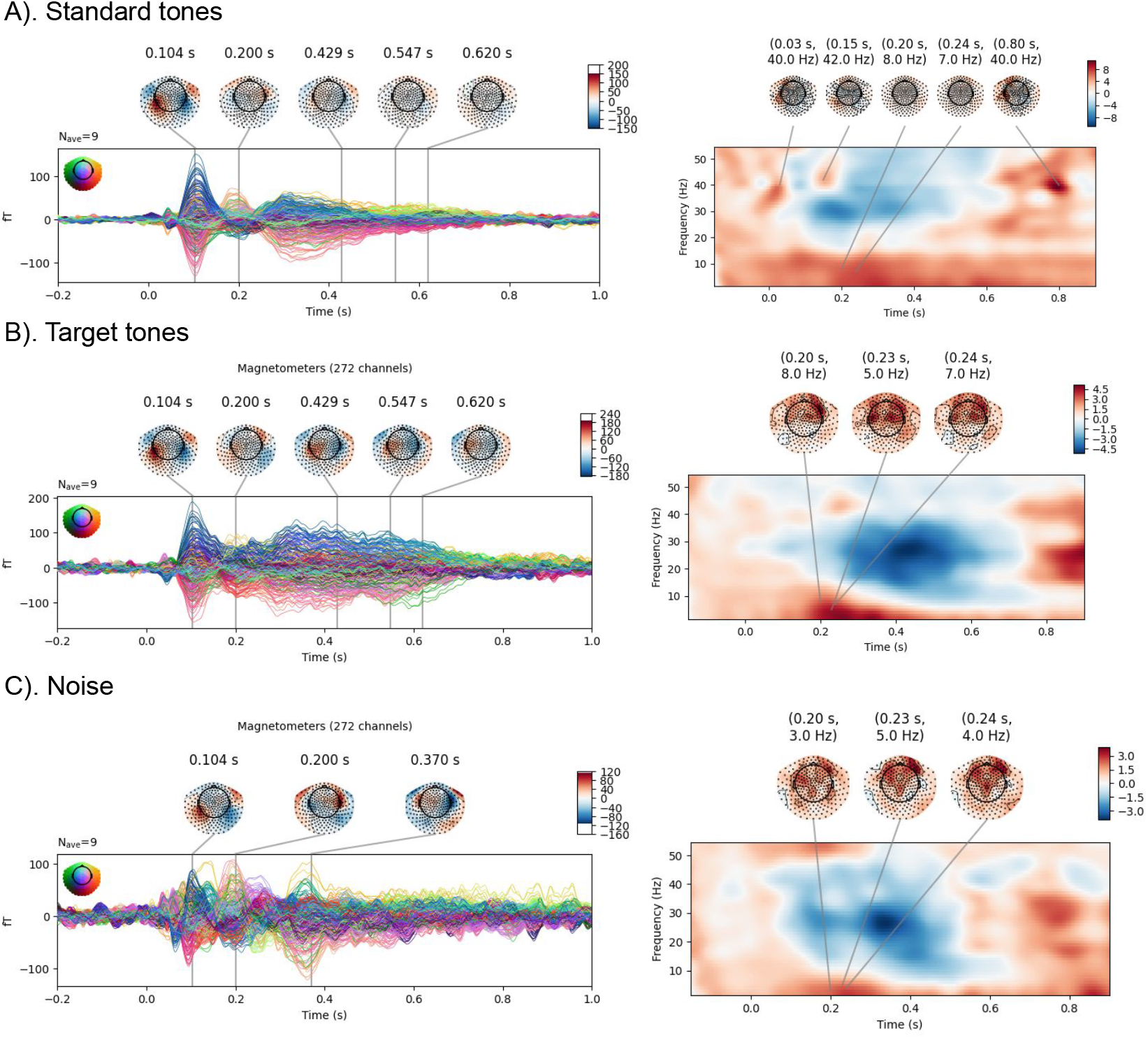
Evoked and induced activity in response to (A) standard tones, (B) target tones and (C) the noise. The color-coded values representing induced activity correspond to z-scores

The MNE-based source localization of the induced activity identified the involvement of the fronto-temporal regions for the theta band(see Fig. 10, A) and bilateral frontal regions for the gamma band (see Fig. 10, B). These individual cortical activation maps were subsequently used to construct the CD-PSIICOS projectors.

**Figure 10:**
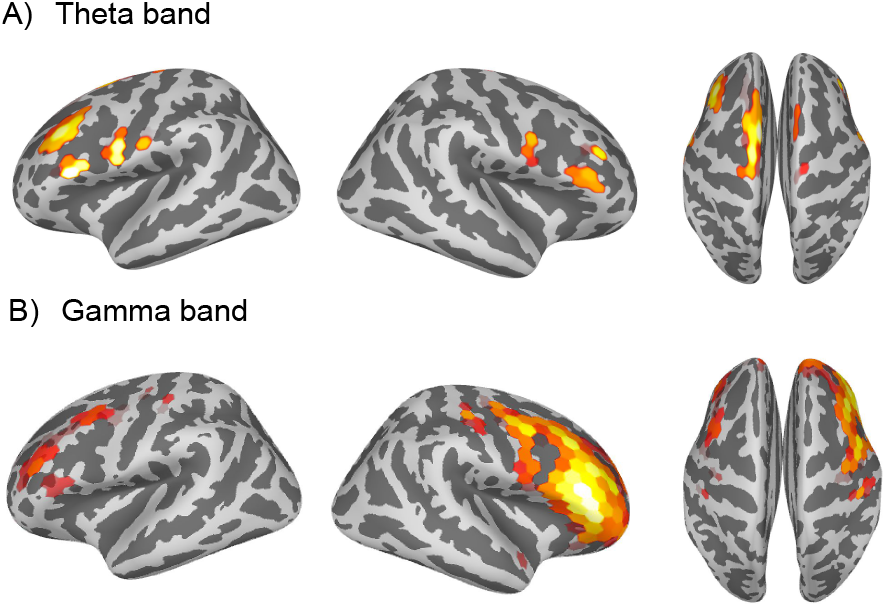
Power profiles of activity in theta (A) and gamma (B) bands obtained with MNE inverse operator and used for construction of CD-PSIICOS projectors

The data-driven attenuation curves, illustrating the balance between suppressing the real part of the cross-spectrum and spatial leakage (SL), showed similar patterns to those observed in the simulations (see Fig.7). The optimal rank for the CD-PSIICOS projector was found to be lower than the optimal rank for the standard PSIICOS (see Fig. 11). For the CD-PSIICOS solutions in the theta and gamma bands, the optimal rank was 40. In contrast, the optimal rank for the standard PSIICOS solutions was equal to 70 for the theta band and 100 – for the gamma band.

**Figure 11:**
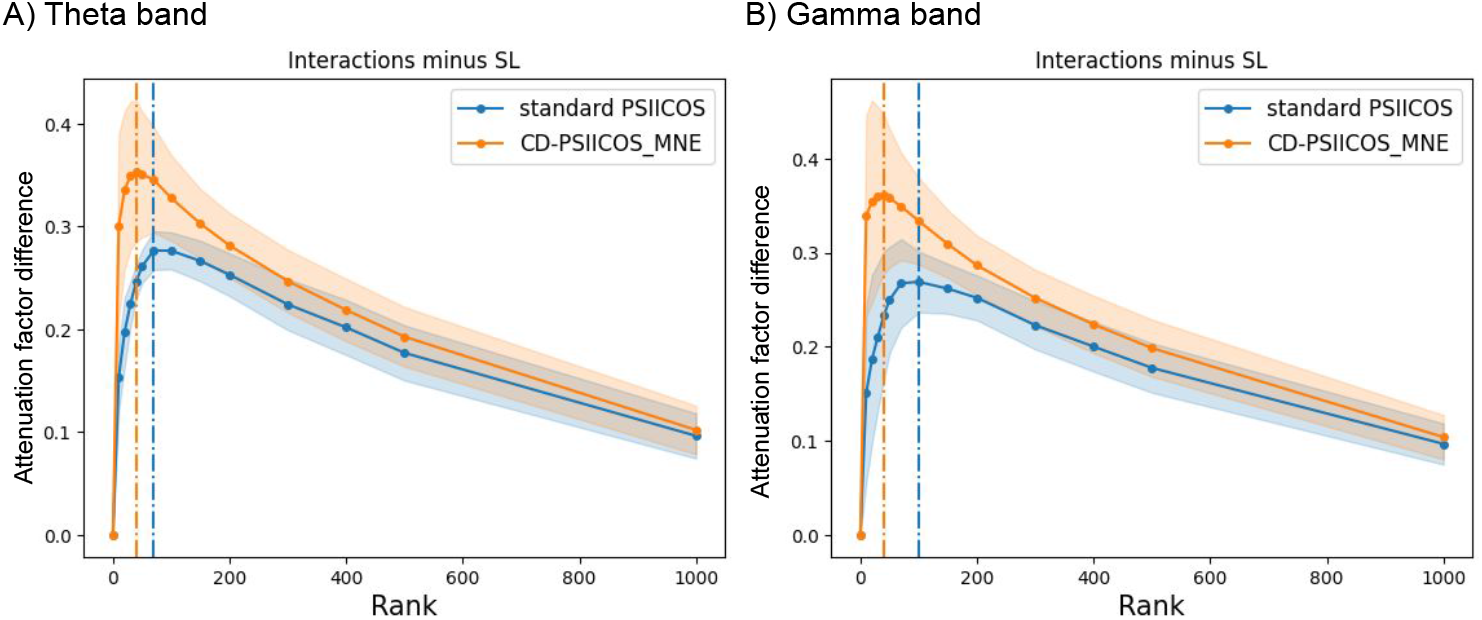
Data-driven attenuation curves for theta (A) and gamma (B) bands. The dashed lines represent the optimal projection ranks. The confidence intervals depict *±SD*.

Additionally, we validated that the observed cross-spectral activity indeed corresponded to the coupling with a consistent phase difference profile, irrespective of the absolute phase values. To this end, we identified the latency of the maximal source-space cross-spectral response (obtained based on the real part of the projected cross-spectrum). Based on this latency, for each trial, we extracted the absolute phases and phase differences for the top most strongly coupled source pairs. Then we created histograms that showed the absolute phases were more uniformly distributed, while the phase differences histograms had peaks around *ϕ* = 0 (see Fig. 12). This result indicates the presence of true phase coupling that cannot be explained by the phase-locked activity.

**Figure 12:**
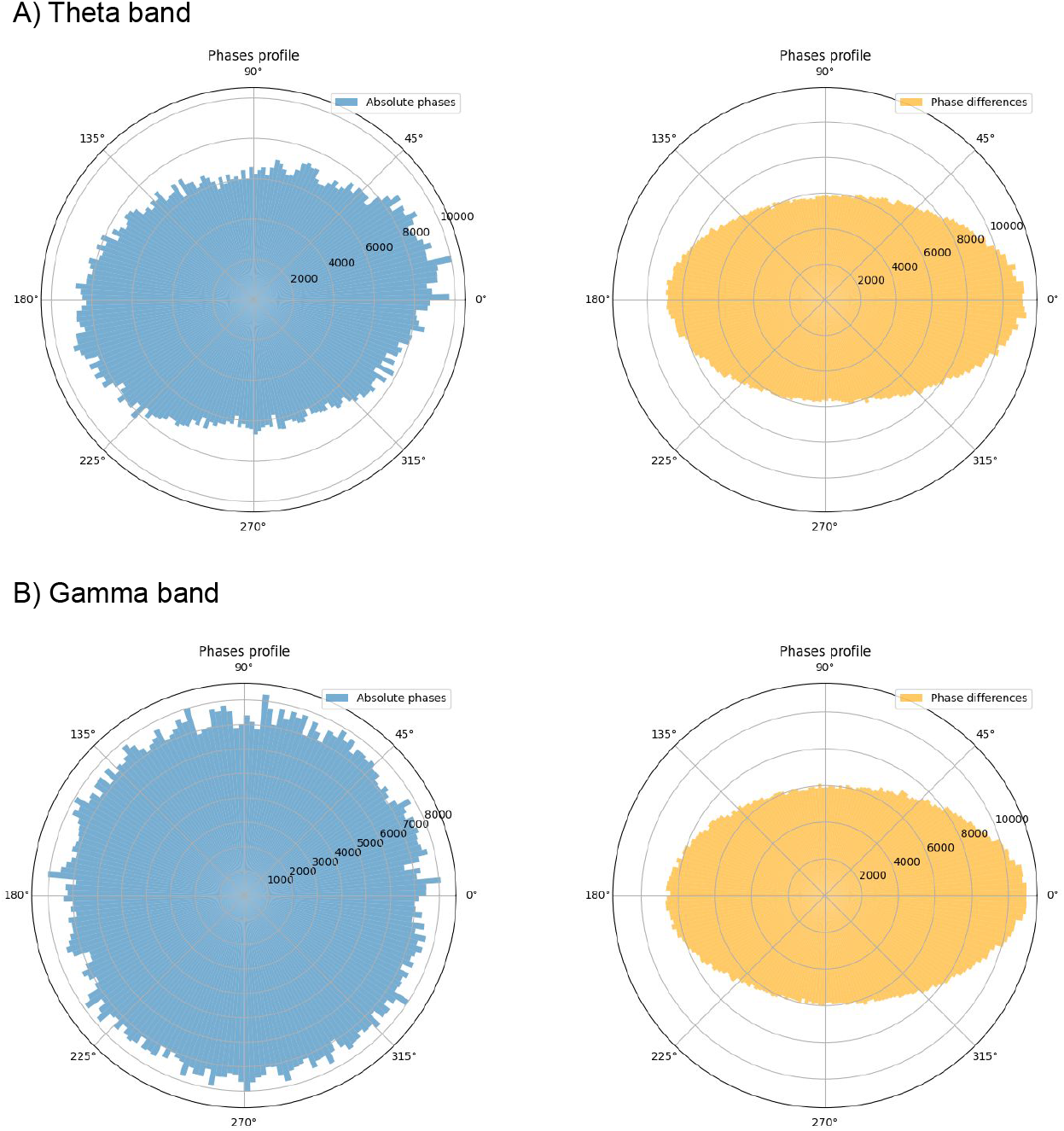
Phase profiles for networks revealed by CD-PSIICOS solution

Finally, we examined the spatial properties of networks and their activation profiles depending on the type of the inverse solution and the projection rank. All the following pictures demonstrate the results of the group analysis over 9 participants.

For the theta band (see Fig. 13), without applying projection, the solution primarily highlights clusters resembling the power distribution of the induced activity. At the projection rank of 10, the standard PSIICOS method shows a partial reduction in spatial leakage but remains biased towards regions with strong power, specifically revealing a distal network with nodes in the left central and right frontal regions. In contrast, the CD-PSIICOS solution at this rank fully suppresses the power-related networks and uncovers a bilateral prefrontal network, characterized by greater spatial symmetry. Additionally, CD-PSIICOS identifies a similar network in the left central and right frontal regions, demonstrating its ability to reconstruct networks comparable to those observed with the standard method. As the projection rank increases to 70, both methods converge on the stable bilateral frontal networks.

**Figure 13:**
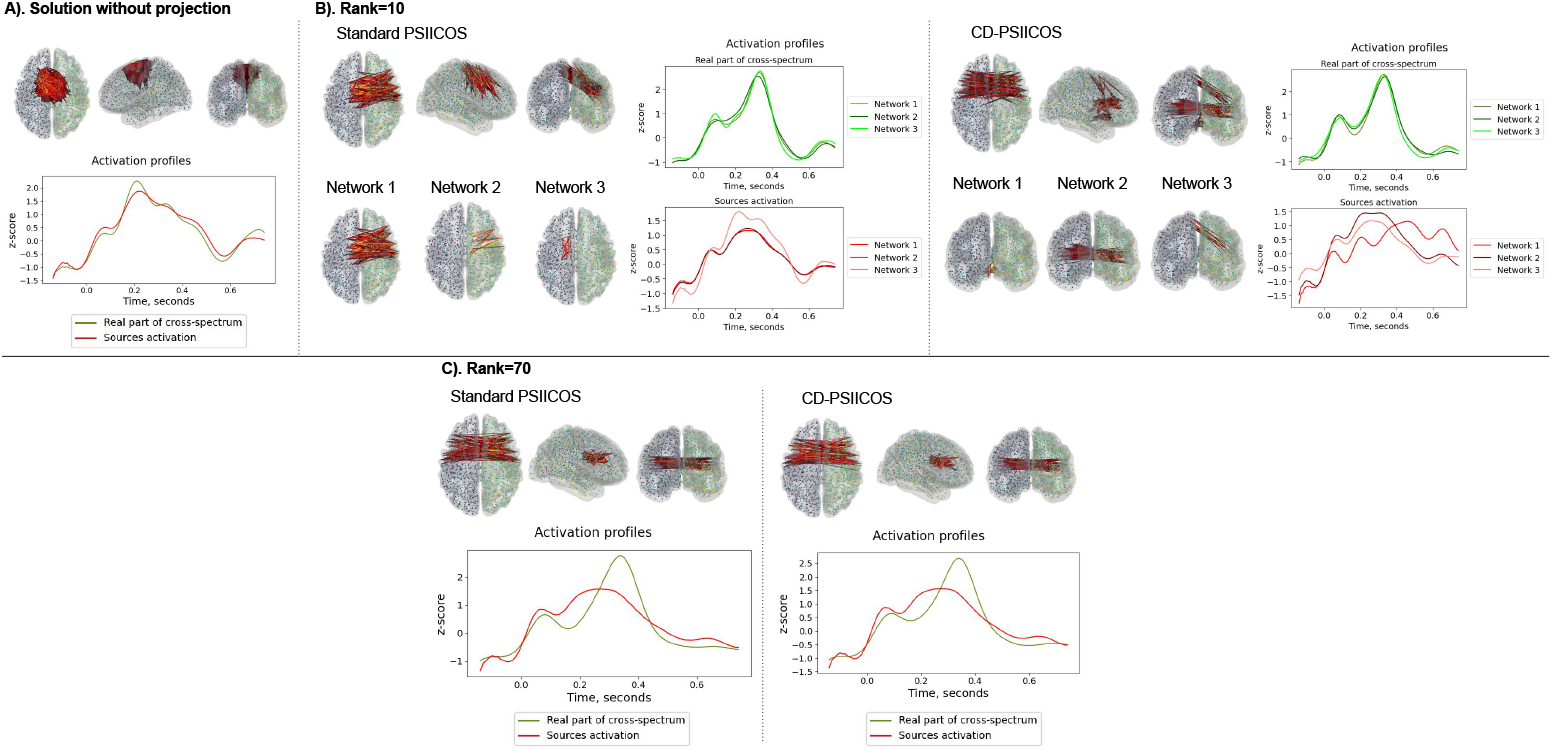
Spatial and activation profiles of the networks in theta band reconstructed without PSIICOS approach (A) and with standard PSIICOS and CD-PSIICOS approaches at ranks equal to 10 (B), and 70 (C). Displayed source pairs correspond to cross-spectral coefficients exceeding 80% of the maximum value

To further analyze the spatial properties of the networks across projection ranks, we identified networks by clustering the spatial patterns of reconstructed connectivity using the KMeans algorithm. In this analysis, each network was defined as the centroid of a spatial cluster derived from the connectivity patterns reconstructed at each projection rank. This approach allowed us to track the persistence of networks as the projection rank increased.

For the theta band (see Fig. 14), in the standard PSIICOS solution, the relevant network (Network 4, Fig. 14, 1.A ) was revealed at higher projection ranks (from the rank of 20). Moreover, at the higher ranks (from 400) the spatial variability of the networks was more pronounced, suggesting reduced robustness in the reconstruction process. In contrast, the CD-PSIICOS solution revealed the stable network (Network 3, Fig. 14, 2.A) at the lower projection rank of 10. Networks identified by CD-PSIICOS remained consistent in their spatial profiles and locations as the rank increased (up to 500).

**Figure 14:**
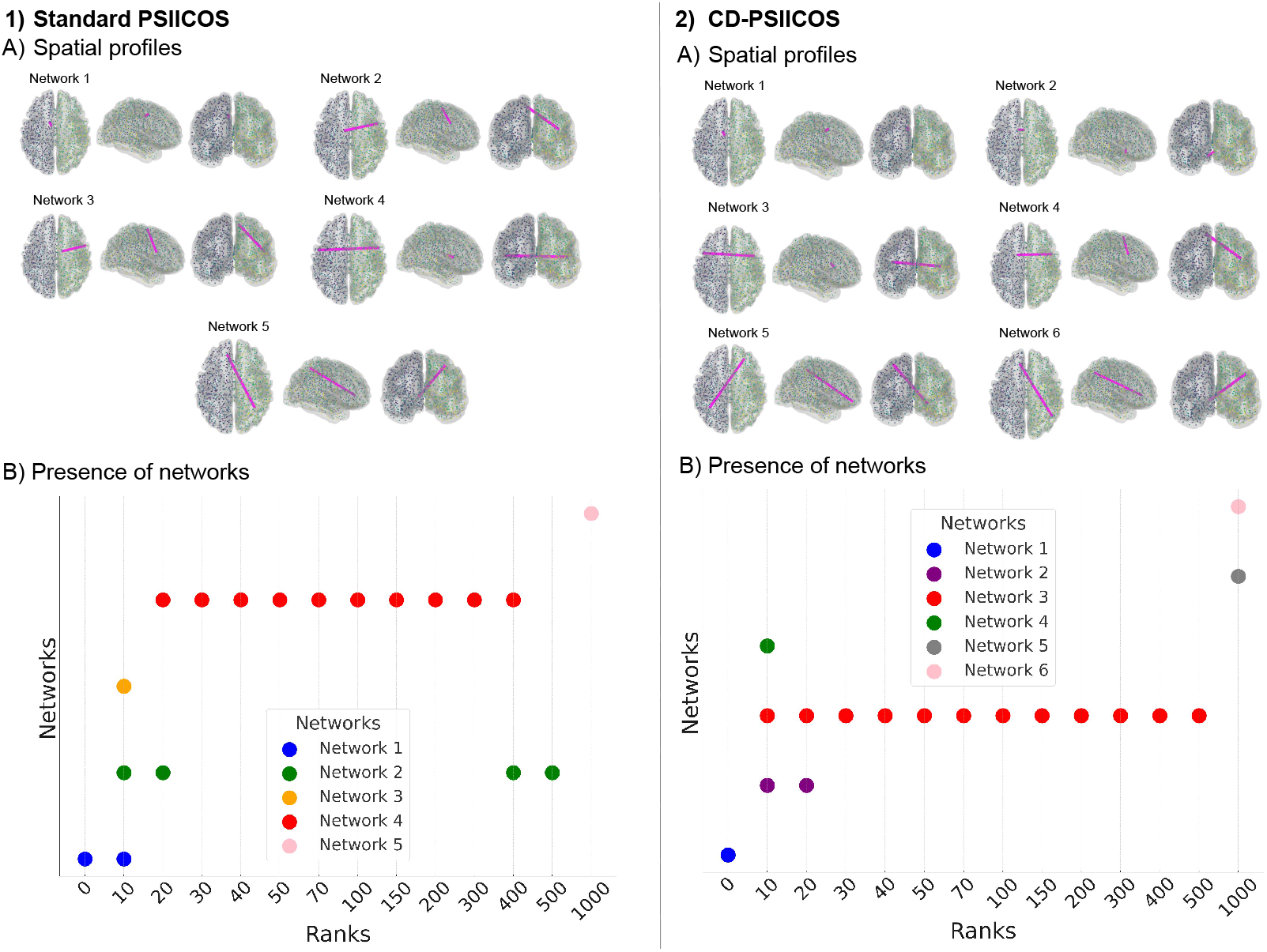
The presence of the networks in the obtained solutions in the theta band depending on the type of projector and the projection rank. The blue, red, green and pink colors indicate the common networks observed in both solutions

In the gamma band, the solution without projection revealed local frontal connections in the right hemisphere (see Fig.15). The standard solution at lower ranks remained biased towards these local connections. However, the CD-PSIICOS solution uncovered more distal networks within the frontal regions.

**Figure 15:**
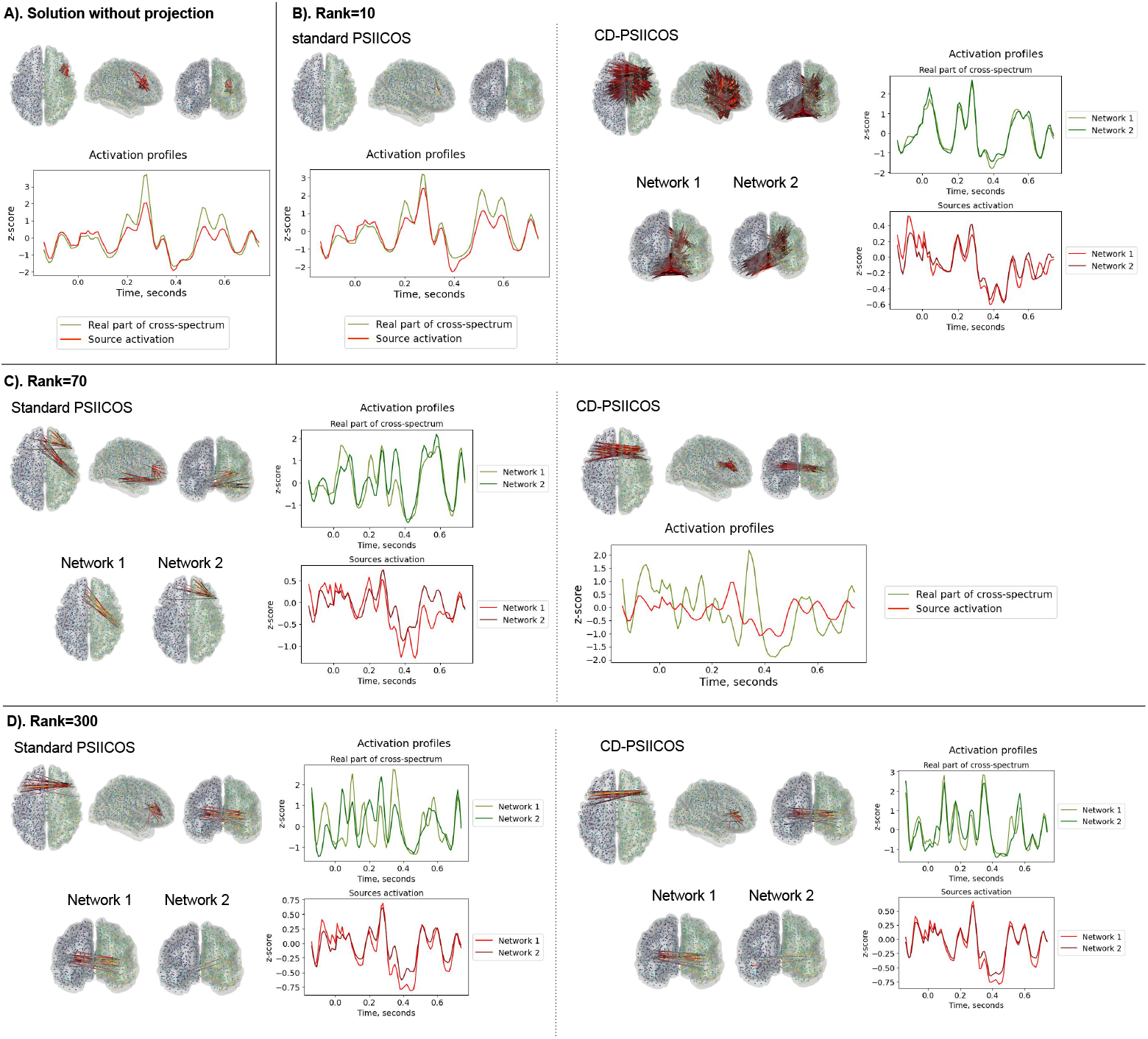
Spatial and activation profiles of the networks in gamma band reconstructed without PSIICOS approach (A) and with standard PSIICOS and CD-PSIICOS approaches at ranks equal to 10 (B), 70 (C), and 300 (D). Displayed source pairs correspond to cross-spectral coefficients exceeding 80% of the maximum value

The standard PSIICOS solution remained biased towards local power-related connections up to the rank of 70 (see Fig. 16), while CD-PSIICOS fully eliminated these connections much earlier at the rank of 10. Starting from the higher ranks of 50 (for the CD-PSIICOS) and 100 (for the standard PSIICOS), both solutions revealed the connections between the left temporal pole and the right frontal and temporal regions (Network 5, Fig. 16, 1.A; Network 3, Fig. 16, 2.A) as well as the frontal bilateral network (Network 4, Fig. 16, 1.A; Network 4, Fig. 16, 2.A). The presence of these networks persisted solution up to the rank of 500. However, at rank 1000, the suppression was so strong that not only were the artifacts eliminated, but the true cross-spectral components of the network were also excessively attenuated, resulting in a loss of meaningful connectivity information and the presence of the long-range fronto-occipital connections.

**Figure 16:**
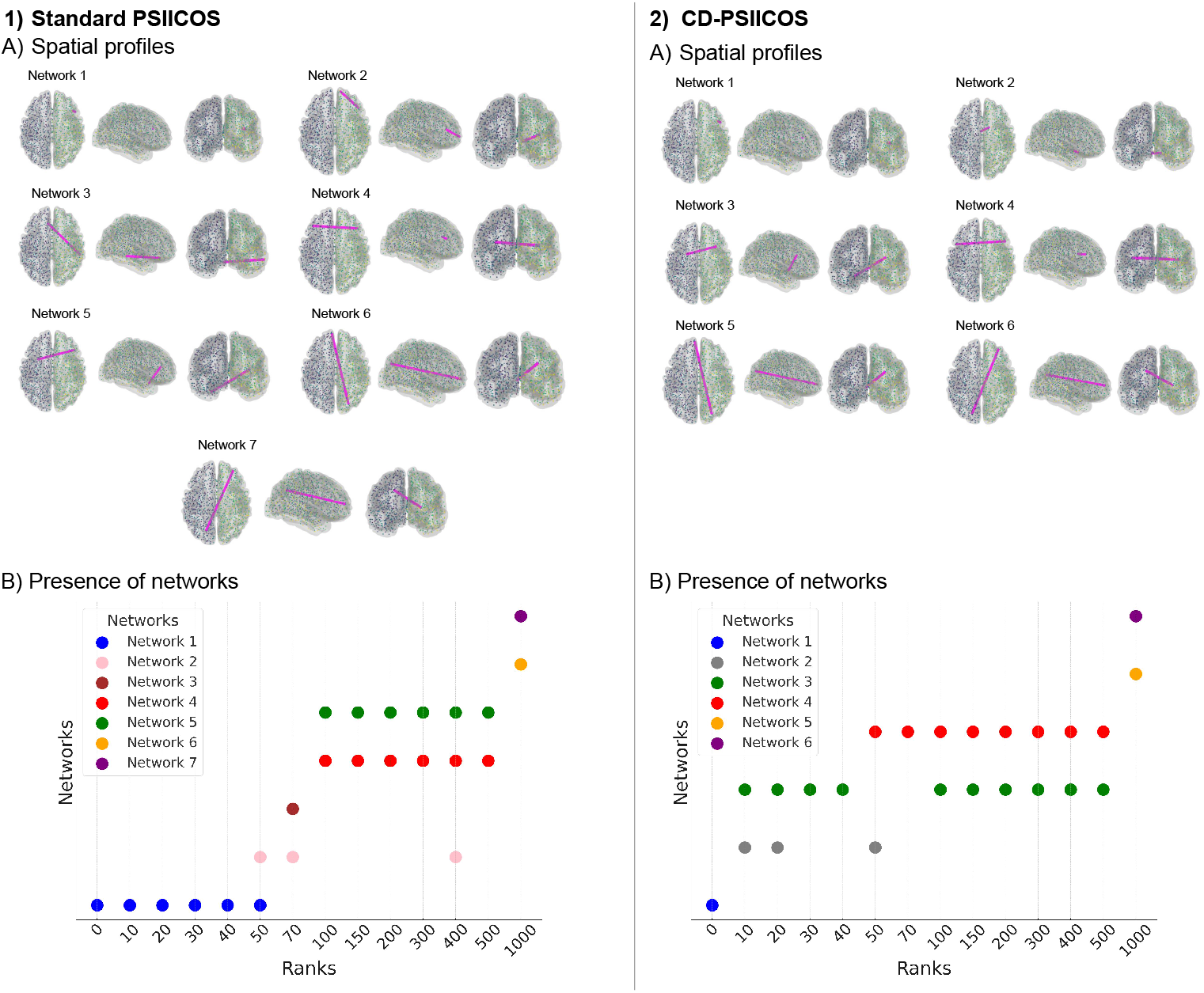
The presence of networks in the obtained solutions in gamma band depending on the type of projector and the projection rank. The blue, orange and green colors indicate the common networks observed in both solutions

The results from the real data demonstrate the improvement of SL suppression obtained by the means of CD-PSIICOS particularly evident in the early suppression of spurious connections and the consistent reconstruction of stable network profiles across ranks. These findings highlight the robustness and reliability of CD-PSIICOS in isolating true functional connectivity, paving the way for more accurate and interpretable network analyses.

## 4. Discussion

This study introduces Context-Dependent PSIICOS (CD-PSIICOS), a novel approach that improves phase-shift invariant estimation of functional networks from MEG/EEG data. Compared to the original PSIICOS technique (Ossadtchi et al., 2018), the CD-PSIICOS more efficiently distinguishes spatial leakage from genuine functional coupling by focusing its suppression on active sources. This enhances the SNR of true coupling components in the projected cross-spectrum, while maintaining the phase-shift invariance property.

The key distinction of CD-PSIICOS from its predecessor lies in the use of the cortical source distribution *p*_*i*_, *i* = 1, … , *N* , to scale the auto 2-topographies when constructing the CD-PSIICOS projector (see equations (7) and (8)). Then as in the original PSIICOS this projector is applied to the vectorized cross-spectrum timeseries matrix to suppress spatial leakage. Subsequently, the matrix is analyzed for components aligned with the interaction 2-topographies **q**_*ij*_, and the corresponding source-space cross-spectral coefficients *c*_*ij*_ are estimated.

Estimating cortical source distribution from EEG or MEG data involves solving the inverse problem, which requires regularization to obtain a unique and stable solution. A key advantage of CD-PSIICOS is its robustness to the choice of regularization parameters in the inverse solvers, greatly enhancing its practical applicability. This contrasts with the traditional functional connectivity (FC) estimation methodology that relies on the inverse problem to directly estimate source time series to be used for computing pair-wise FC statistics such as coherence, phase locking value, etc. This traditional framework is highly sensitive to the regularization parameter of the inverse solver (Vallarino et al., 2023; Hincapié et al., 2016), a limitation that CD-PSIICOS efficiently overcomes (see Figure 8.C).

Our previous work demonstrated the advantage of PSIICOS over several modern FC estimation methods, particularly for cortical sites pairs with activity exhibiting consistent zero or near-zero mutual delays. In this study, realistic simulations indicate that CD-PSIICOS consistently outperforms the standard PSIICOS approach. It also exhibits robust performance across various inverse solutions—including MNE, sLORETA, and DICS—with MNE-based weighting providing a modest but consistent benefit. Furthermore, data-driven attenuation curves reveal that CD-PSIICOS achieves optimal performance at significantly lower projection ranks than the standard PSIICOS method, see Figures 6,7,11, suggesting more efficient and targeted suppression of spatial leakage while preserving genuine functional interactions.

CD-PSIICOS also appears to mitigate the important concerns raised by Schoffelen and Gross (2009) about erroneous functional connectivity (FC) due to amplitude changes in sources and due to signal-to-noise ratio (SNR) alterations from surrounding noise (Muthukumaraswamy and Singh, 2011). To demonstrate this, in our simulations we introduced a pair of uncoupled sources with amplitudes three times greater than those of the four other sources forming the true network nodes. As shown in Figure 5, CD-PSIICOS accurately reconstructed the ground-truth FC configuration at a low projection rank of R=80, and the solution remained stable as the rank increased. Conversely, the original PSIICOS approach required a higher rank of R=200 for comparable performance, potentially suppressing more subtle but genuine interactions.

Validation using real MEG data from nine healthy participants further corroborated the efficacy of CD-PSIICOS. The results align with established findings on the involvement of theta oscillations in auditory discrimination and target detection, particularly in mismatch- and P3-related dynamics (Harper et al., 2017; Polich, 2007; Ko et al., 2012). Observed theta coupling between lateral prefrontal regions in both hemispheres supports the role of these oscillations in goal-directed behavior, cognitive control, and sustained attention (Rajan et al., 2019; Christian et al., 2023). Furthermore, gamma-band frontal networks are consistent with activation of the ventral attention system, implicated in bottom-up attentional capture and distractor processing (ElShafei et al., 2020). CD-PSIICOS consistently identified physiologically plausible functional connections at lower projection ranks than the original PSIICOS, demonstrating superior suppression of spatial leakage and task-related power artifacts while preserving true connectivity estimates. Notably, CD-PSIICOS exhibited greater stability and consistency across projection ranks, as evidenced by the persistence of meaningful networks and the absence of spurious or artifactual connections.

Both PSIICOS and CD-PSIICOS rely on the estimation of the unnormalized source-space cross-spectral coefficient 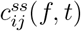, whose magnitude reflects contributions from both the instantaneous amplitude correlation and the mutual phase consistency of rhythmic activity between neuronal populations. The relative independence of amplitude and phase coupling mechanisms (Siems and Siegel, 2020; Bruns et al., 2000) complicates the interpretation of the specific type of coupling identified by either method within the PSIICOS family. While our simulations demonstrated that CD-PSIICOS remains robust against spurious influences from active but uncoupled sources, the reliance on an un-normalized metric highlights the need for the development of appropriate statistical procedures. In particular, establishing confidence intervals for the estimated source-space cross-spectral coefficients would provide a principled framework for assessing the significance and reliability of the detected interactions.

Most functional connectivity (FC) detection methods assume the presence of pronounced oscillatory activity at each node being tested for synchrony. However, an increase in the cross-spectral coefficient 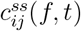 can still occur due to enhanced synchrony between sources *i* and *j*, even if it coincides with local desynchronization at one or both nodes—leading to a reduction of oscillatory power at the corresponding locations. Importantly, as long as remote coupling persists, it may remain detectable by subspace-tracking approaches such as PSIICOS and CD-PSIICOS. Investigating the performance of these methods under conditions of local desynchronization represents an important and promising direction for future research. This could be explored initially through appropriately designed realistic simulations and subsequently validated in well-characterized experimental paradigms, such as the desynchronization of sensory-motor rhythms (alpha and beta bands) induced by motor imagery or actual movement execution.

In conclusion, CD-PSIICOS represents a refinement in functional connectivity analysis by incorporating task-specific contextual information in the form of source power distribution. By incorporating data-driven weights derived from power distributions, the projector effectively focuses on relevant source-space interactions while suppressing spatial leakage and spurious connections. This approach not only enhances the accuracy of connectivity estimates but also extends the framework for future studies that demand precise differentiation between genuine neural interactions and task-related power modulations, especially in relation to ubiquitous in the brain zero or close-to-zero phase-lag couplings.

## 7. Code and data availability

The dataset with the real MEG recordings from the healthy participants (Nugent et al., 2022) can be accessed via OpenNeuro (doi:10.18112/openneuro.ds005752.v2.1.0).

## 8. Ethics statement

This study did not record any new dataset neither from animal nor from human subjects. The study used a preexisting MEG dataset recorded from healthy participants and published in (Nugent et al., 2022) along with the necessary ethics statements.

